# Effect of the daily exposure of 1.7 GHz LTE electromagnetic field in early fetal brain development using a human brain organoid model of the cerebral cortex

**DOI:** 10.1101/2024.11.20.624497

**Authors:** Gyu Yeon Park, Ya Qi Pan, Yoon-Jeong Choi, Nam Kim, Sangbong Jeon, Kiwon Song

**Author notes:** Address all correspondence to Kiwon Song, Department of Biochemistry, College of Life Science and Biotechnology, Yonsei University, Seoul 03722, Korea.

## Abstract

Wireless communication technology’s ubiquity means constant exposure to radiofrequency electromagnetic fields (RF-EMF). While concerns exist about RF-EMF’s effects on fetal brain development, studying this in pregnant women poses ethical and safety challenges. Our study examined 1.7 GHz LTE RF-EMF’s impact on early human fetal brain development using brain organoids derived from human induced pluripotent stem cells (hiPSCs).

Previous research showed that 100-day-old human cerebral cortical brain organoids correspond to 19-24 weeks post-conception of fetal brain development. We exposed organoids to 1.7 GHz LTE RF-EMF at 8 W/kg specific absorption rate (SAR) for 5 hours daily up to 100 days. We monitored size, developmental morphology, and developmental stage markers using immunofluorescence microscopy and qPCR analysis.

Comparing RF-EMF-exposed organoids with controls showed no significant differences in size, development, or expression of key markers: SOX2 and PAX6 (pluripotent cell markers), NEUN (mature neuronal marker), MAP2 (dendrite marker), and SYN1 (synapse marker). Results suggest that daily exposure to 1.7 GHz LTE RF-EMF at 8 W/kg SAR may not significantly affect early human fetal brain development.

## 1. Introduction

With the development of telecommunication technologies, wireless devices are widely used in daily life. As the usage of mobile phones increases, humans are consistently exposed to radiofrequency electromagnetic fields (RF-EMFs). RF-EMF within the frequency range of 100 kHz to 300 GHz is utilized for long-distance transmission in wireless communication [3]. As long-term evolution (LTE) technologies have enabled the convergence of wired and wireless networks such as GSM, LAN, and Bluetooth [3–5], LTE is currently the most widely adopted telecommunication technology. LTE technology in the 1.7 to 1.8 GHz range is predominantly used in 4^th^ generation mobile technology in Korea [6–9]. Thus, there are increasing concerns about the effect of RF-EMF generated by 1.7 GHz LTE on the human body. According to the International Commission on Non-Ionizing Radiation Protection (ICNIRP), the safety limits for electromagnetic radiation from mobile devices for the general public are specified as 4 W/kg for the limbs, 2 W/kg for the local head/torso, and 0.08 W/kg for the entire body. However, the physiological effects of 1.7 GHz LTE RF-EMFs on humans remain controversial [3, 10].

The World Health Organization (WHO) has described RF-EMF as one of the most critical environmental issues and established *The International EMF Project* to define safety standards for EMF in the frequency range from 10 MHz to 300 GHz [4, 5]. Within the 10 MHz-300 GHz RF-EMF range of the International EMF Project, effects on pregnancy and birth outcomes were regarded as one of the critical issues [11, 12]. During pregnancy, the fetus in early developmental stages can be highly sensitive to environmental influences, including RF-EMF, increasing the risk of abnormal growth and development [13]. Especially, the effect of RF-EMF on the brain development of early fetuses has been a serious concern of the public, and several studies have been performed with model mammalian organisms. Chen *et al.* demonstrated that 4 W/kg 1800 MHz RF-EMF exposure on embryonic neural stem cells from embryonic day 13.5 mouse cortex did not alter the proliferation and the ratio of differentiated neurons and astrocytes but decreased expression of the proneural genes Ngn1 and NeuroD, which are crucial for neurite outgrowth [14]. Saghezchi *et al.* also showed a negative effect on the development of osteogenesis when exposing the forelimbs of prenatal mice to 2.4 GHz RF-EMF for 4 h/d until delivery [15]. In Koohestanidehaghi *et al*., mice zygotes produced by *in vitro* fertilization showed a decreased cleavage rate and reduced time to reach the blastocyst stage by 30 min exposure to RF-EMF at 900-1800 MHz for 5 days [16]. On the other hand, Shirai *et al.* showed that exposing the whole body of pregnant rats and newborns to 2.14 GHz W-CDMA RF-EMF for 20 h daily at < 0.24 W/kg did not result in any adverse effects [17]. Another study reported that exposure to 2.45 GHz Wi-Fi signals for 2 h per day, 6 days per week for 18 days at 4 W/kg induced no abnormalities in pregnant rats and in the pre- and post-natal development of the fetus [18]. These controversial physiological effects of RF-EMF reported on the fetal development in various mammalian models strongly suggest that additional studies are required to reveal the accurate effects of RF-EMF on human fetal development.

Recently, human organoids have been used as an innovative approach for modeling human tissue and organ development. Cerebral cortical organoids derived from human induced pluripotent cells (hiPSCs) show three-dimensional neural-tissue-like structures resembling the fetal human brain [2]. In human brain organoid formation from hiPSCs, neuroepithelial cells (NEs) in the early stage of organoids (∼ day 20) form a rosette structure in the ventricular zone (VZ), which is a pseudostratified columnar epithelium consisting of actively proliferating NEs and neural stem cells [19, 20]. NEs from neural stem cells then start to transform into apical radial glial cells (aRGs), which serve as the primary type of neural progenitor cells during corticogenesis. These aRGs located in the VZ proliferate to increase the pool of progenitor cells. In the middle stage of organoids, radial glial cells (RGs) generate intermediate progenitor cells (IPCs), which migrate to the subventricular zone (SVZ) to differentiate into deep- and superficial-layer neurons and to construct neural circuits in the late stage of the human brain organoid formation [21, 22]. Synaptic formation, function, and elimination are controlled by astrocytes. Astrocytes have been reported to start developing after day 50, and mature astrocyte markers begin to be detected between day 50 and 100 in human cortical organoids [23]. Additionally, astrocytes secrete essential cytokines for neural and synaptic maturation from early gestation, such as TNFα and IL-1β [24]. Yoon *et al.* demonstrated through transcriptomic analysis that human cortical organoids at day 100 highly overlap with the mid-fetal prenatal brain (19-24 post-conception weeks, PCW) [2]. Moreover, neurons extend axons and dendrites and start synapse connections during 18-22 PCW [25]. Synaptogenesis, a critical developmental process in the nervous system, can be observed in ∼100-day organoids. Thus, hiPSC-induced human brain organoid development for 100 days provides an excellent model system for fetal cerebral cortex development of the human brain.

In this study, we aimed to analyze the physiological effect of 1.7 GHz LTE RF-EMF on human fetal brain development using a hiPSC-induced human brain organoid model of the cerebral cortex. We exposed human brain organoids to 1.7 GHz LTE RF-EMF at a specific absorption rate (SAR) of 8 W/kg daily during their growth and development for 100 days and assessed its effects on early human fetal brain development.

## 2. Results

### 2.1 Human cortical brain organoids from hiPSCs mimicked the fetal brain development

To investigate the effects of RF-EMF on fetal brain development, we first generated human cortical brain organoids from hiPSCs that closely recapitulate the process of fetal brain development. hiPSCs are derived from adult somatic cells by overexpressing Yamanaka factors, including SOX2 and OCT4 [24], which are crucial in maintaining their pluripotency, allowing them to differentiate into various cells in tissues and organs [27]. To ensure the generation of high-quality brain organoids, we validated the pluripotency of ASE9209 iPSCs used in this study through quantitative polymerase chain reaction (qPCR) analysis. Compared with HeLa cells, the expression of OCT4 and SOX2 remained high in ASE9209 culture, indicating the pluripotency of these cells (Figure S1A). We also observed that ASE9209 iPSCs exhibited low expression of the endoderm marker SOX17 and the mesoderm marker BRACH, while highly expressing the ectodermal marker SOX2 (Figure S1). Considering that the brain primarily derives from ectodermal cells of the early developmental stage, we concluded that ASE9209 cells with high ectoderm marker expression would be suitable for brain cortical organoid generation. We also checked the quality of ASE9209 cells by monitoring for mycoplasma infection but could not detect any sign of infection (Supplementary Figure S1C).

We then started to produce human cortical brain organoids with quality-verified ASE9209 cells by following Pasca *et al.* and Yoon *et al*., as described in Materials and Methods [2, 26]. To ensure the growth of brain organoids, their size and shape were monitored for up to 64 days. The size of organoids exhibited a pronounced increase with round shapes during the proliferation and differentiation stages (Figure 1A and Figure S2), as reported by Yoon *et al.* [2]. In parallel, we examined the differentiation process of organoid development to confirm that brain organoids recapitulate human brain architecture by checking the structure and the expression of markers specific to each developmental stage of the neurulation process. During the early developmental stage, brain organoids undergo neurulation and form rosette structures within the VZ. Immunofluorescence microscopy revealed that SOX2 expression was widely scattered around cells in rosette structures on day 25 organoids, which decreased and compacted in the VZ on day 50 organoids (Figure S3A). The differentiation of neural stem cells in the VZ into neurons and their migration into the SVZ were also confirmed by the expression of the neural progenitor cell marker PAX6 and neuron marker NEUN in layers of the day 50 brain organoids (Figure S3B). These results indicate that human brain organoids we generated from ASE9209 cells mimic the fetal brain developmental process and successfully resemble the structure of the forebrain cerebral cortex in the human brain making them suitable for RF-EMF exposure studies.

**Figure 1.**
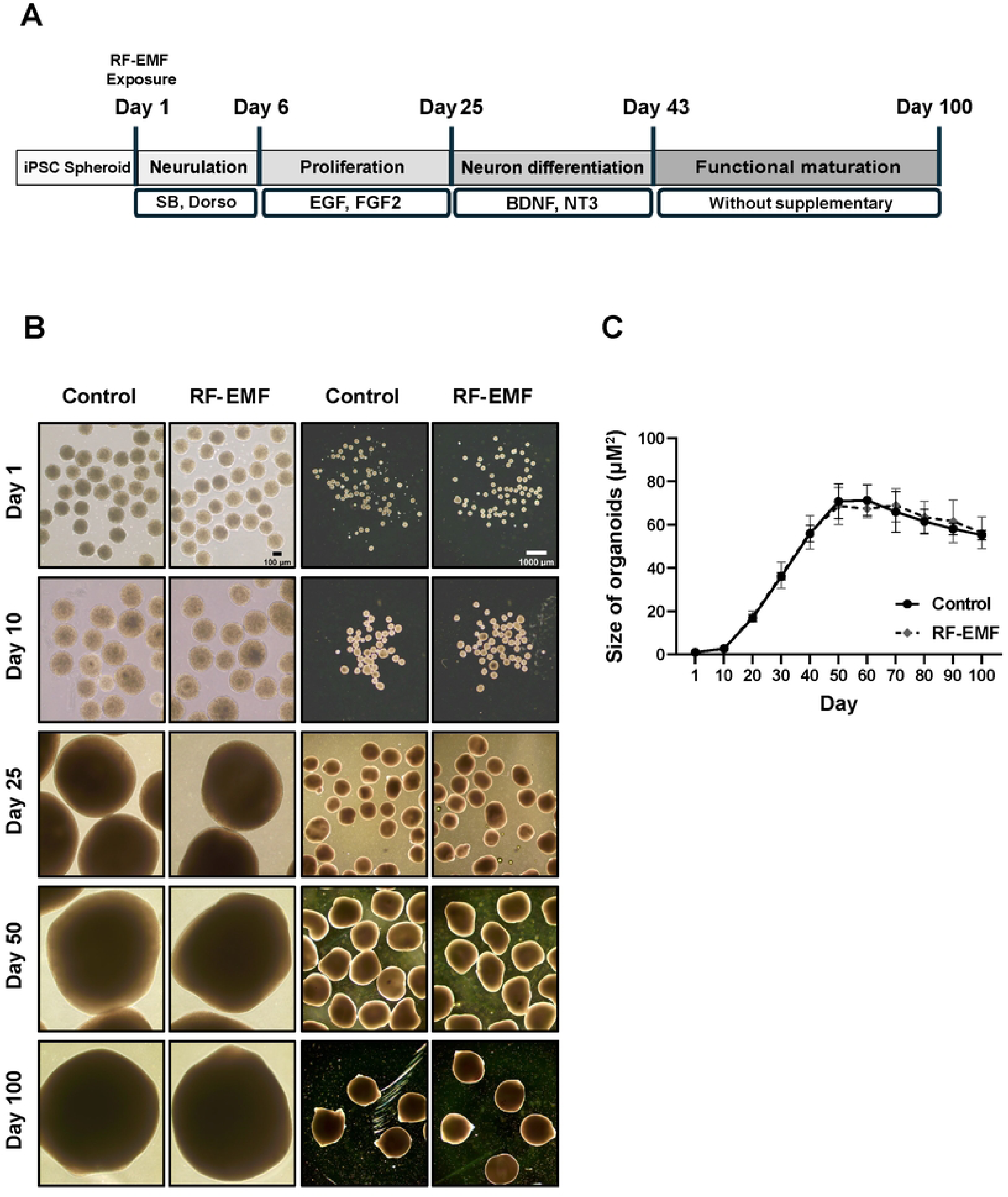
The morphological characteristics of 1.7 GHz LTE RF-EMF-exposed and the sham control human cortical brain organoids. (A) The experimental scheme illustrates the process of hiPSC-derived human brain organoid generation applied in this research. (B, C). The RF-EMF-treated groups of human cortical brain organoids were exposed to 1.7 GHz LTE RF-EMF at SAR of 8 W/kg for 5 h daily for 100 days with sham controls. (B) Representative images of the RF-EMF-exposed and sham control human cortical brain organoids at each designated day of growth. A Nikon microscope (ECLIPSE Ts2) with a 4 X and 1 X objective lens was used to capture these images. Scale bars, 100 μm for Black and 1000 μm for White. (C) A graph represents the size of the RF-EMF-exposed and sham control human cortical brain organoids in 100 days. Approximately 10 to 60 organoids were measured every 10 days for both the sham control and the RF-exposed groups, and the average values were calculated from 3 independent experiments. The average size of the organoids was assessed using the ROI manager in the Image J software. Statistical analysis was conducted using multiple paired t-tests on the data from three independent experiments.

### 2.2 1.7 GHz LTE RF-EMF did not affect the developmental size of human cortical brain organoids

The *in vitro* radiofrequency radiation exposure device used in this study was a radial transmission line (RTL) exposure system that exposed multiple cell plates simultaneously [7–9]. In previous studies, we developed 1.7 GHz LTE RF-EMF-generating device and showed that RF exposure induced decreased cell proliferation and senescence [7, 8]. By upgrading the device with stable power monitoring and efficient thermal control, we showed that the exposure of 1.7 GHz LTE RF-EMF at SAR of 0, 0.4, 1, 4, and 8 W/kg induced no changes in the proliferation of various human cells, compared with the sham exposure controls [9]. In this study, to assess the physiological effect of 1.7 GHz LTE RF-EMF on human brain organoids, we used the same RF-EMF generating device employed in the previous studies [7–9]. The schematics of RF-EMF system applied were shown in the Supplementary Figure S5 and the detailed information on the device was described in Choi *et al*. [7], and briefly in Materials and Methods. We exposed developing human cortical brain organoids to 1.7 GHz LTE at the SAR of 8 W/kg for 5 h daily for 100 days. The SAR value was determined based on the input power (W) as shown in Table 1. We chose 8 W/kg as the SAR value of exposure since it is the highest RF-EMF strength that does not generate heat in our device, and exposure to a SAR of 8 W/kg did not show any effect on human cell proliferation [9]. If the exposure to a SAR of 8 W/kg does not induce any effect on human brain organoids, we could infer no effect with SAR values lower than 8 W/kg. We decided on RF-EMF exposure time of 5 h daily, considering the report that the average exposure time to cell phones for American adults was 5 h/day [29].

**Table 1.**
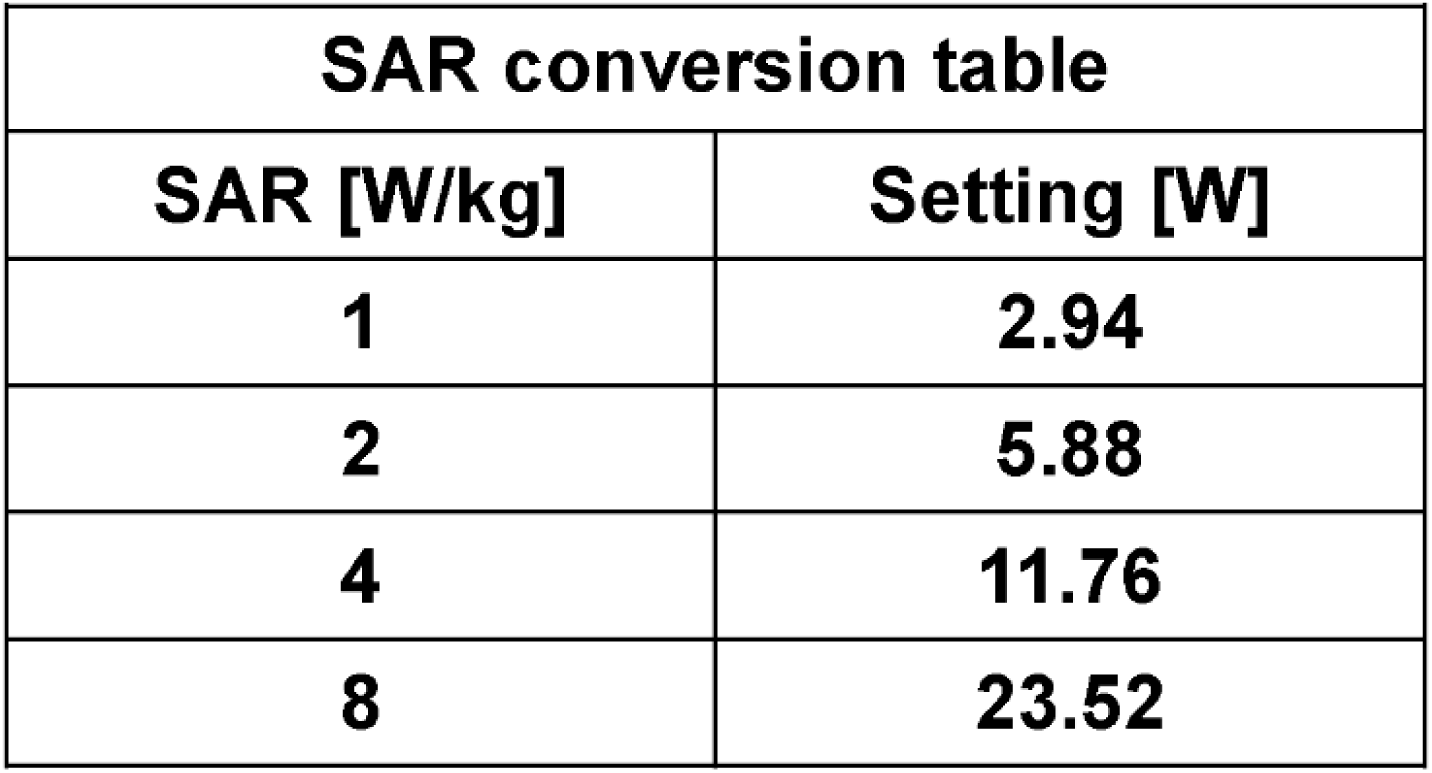
Conversion table for specific absorption rate (SAR)

| **SAR conversion table** |  |
| --- | --- |
| **SAR [W/kg]** | **Setting [W]** |
| **1** | **2.94** |
| **2** | **5.88** |
| **4** | **11.76** |
| **8** | **23.52** |

Relatively uniform spheroids from ASE9209 were generated using Aggrewell^TM^800 and transferred to Neurulation media for organoid development to start RF-EMF exposure (Day 1 in Figure 1A). The shape and size of 1.7 GHz LTE RF-EMF exposed organoids were monitored daily for 100 days using microscopy and compared with those of the sham control group. As shown in Figures 1B and C, no significant difference in either the shape or the size between the RF-EMF-exposed and the sham group was observed. Hence, these observations suggest that daily exposure to 1.7 GHz RF-EMF does not cause obvious effects on the development of human brain organoids in terms of size and morphology.

### 2.3 1.7 GHz LTE RF-EMF did not affect the formation of ventricular zone in early human cortical brain organoids

In the early stage of brain organoid development, hiPSC spheroids undergo neurulation to differentiate into neural progenitor cells. Around day 15-20, neural progenitor cells can be observed in a rosette structure within the VZ, and their numbers gradually decrease as they differentiate into neural cells [30, 31]. To investigate whether the exposure to 1.7 GHz LTE RF-EMF affects neurulation in the early stage of brain organoid development, organoids were sectioned and stained for immunofluorescence microscopy with the progenitor stem cell marker SOX2 at days 25, 50, and 75. In both the RF-EMF-exposed organoids and the sham control, we observed that progenitor cells were present in VZs (Figure 2A, white arrows) and differentiated neuronal cells extended outside of the VZ (Figure 2A, red arrows). SOX2 was distributed throughout in the progenitor cells of the RF-EMF-exposed-day 25 organoids as in the sham controls (Figure 2A). As development progressed, SOX2 expression was gradually reduced and became concentrated around the VZ in RF-EMF-exposed day 50 and day 75 organoids as in the sham controls (Figure 2A), which is consistent with the observations shown in Figure S3A.

**Figure 2.**
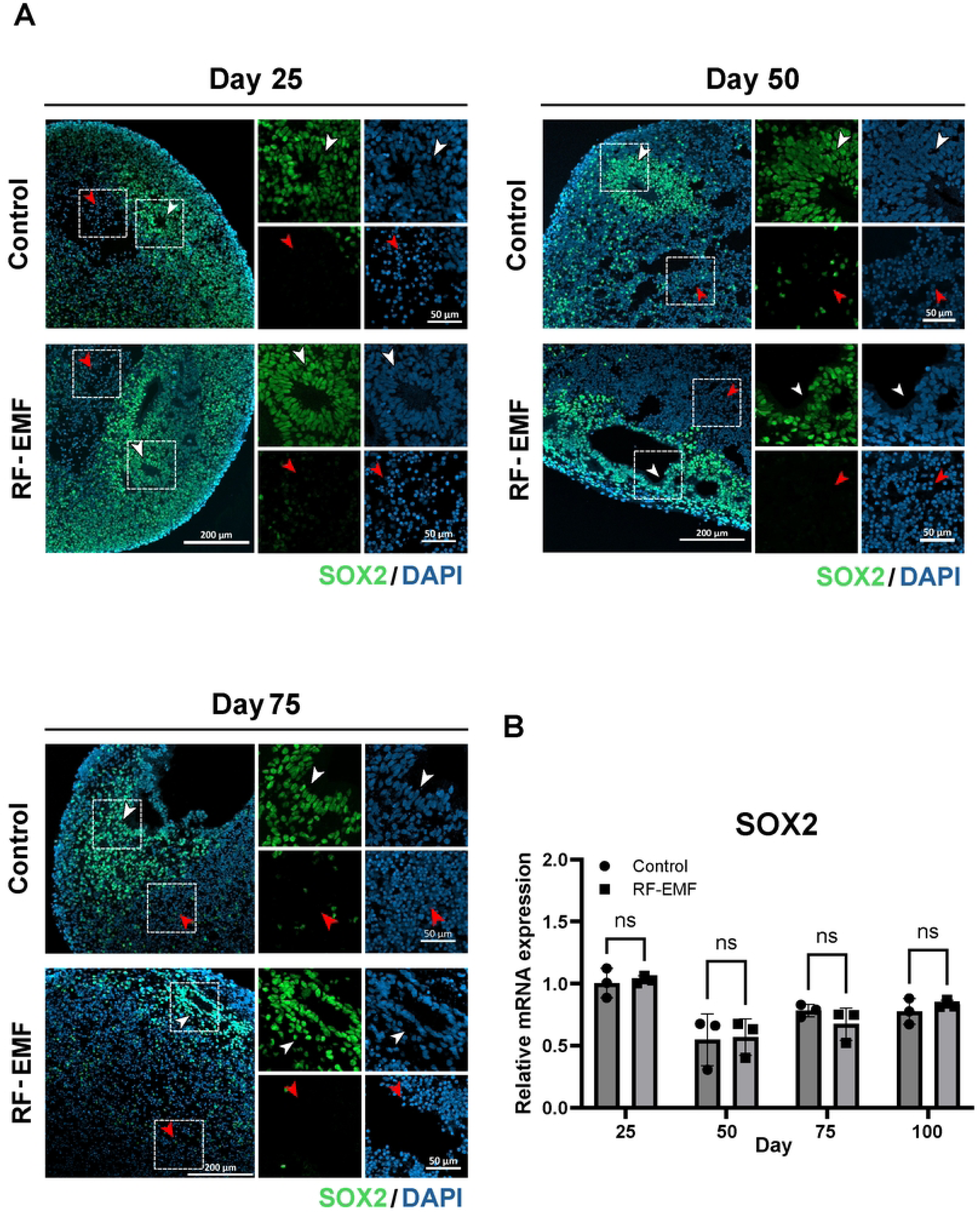
The neural progenitor cells in the developmental process of the human cortical brain organoids exposed to 1.7 GHz LTE RF-EMF and the sham controls. The developing organoids were exposed to 1.7 GHz LTE RF-EMF at a SAR of 8 W/kg for 5 h every day with the sham controls. (A) Organoids were sampled at 3 time points of differentiation (days 25, 50, 75) for immunofluorescence microscopy. The expression of SOX2 (Neural progenitor cell) was observed in VZ surrounding a lumen, illustrating differences in cell morphology between the progenitor cells (white arrow) and the differentiated cells (red arrow). Images were captured using a LSM900 confocal microscopy (Carl Zeiss) with detectors (GaAsP-PMT 2ea, additional GaAsP-PMT 1ea), using 10X and 20X objectives. Scale bar, 200 μm (left) and 50 μm (right). (B) The mRNA expression level of SOX2 in developing organoids was analyzed using qPCR of developing organoids (n=3) at 4 time points (days 25, 50, 75, 100). qPCRs were performed in triplicate for each of the 3 independent experiments and the data were analyzed using multiple paired t-tests of the Holm-Sidak method. The expression levels of SOX2 were all normalized using the expression of actin as a reference gene. The relative mRNA expressions of normalized SOX2 were plotted over the average SOX2 expression of the day 25 sham control organoids. The graphs represent the average values ± SEM. P > 0.05 was considered statistically not significant, ns.

We then confirmed the relative mRNA expression of SOX2 by qPCR in the day 25, 50, 75, and 100 RF-EMF-exposed organoids and sham controls. As shown in the graph of Figure 2B, the expression of SOX2 was at the maximum level on day 25 and decreased on days 50, 75, and 100 in both the RF-EMF-exposed organoids and the sham controls, demonstrating no significant difference in SOX2 expression between the two groups (Figure 2B). These observations indicate that 1.7 GHz LTE RF-EMF at a SAR of 8 W/kg did not affect the formation of VZ and neurulation of neural progenitor cells during the early stage of brain organoid development.

### 2.4 1.7 GHz LTE RF-EMF exposure did not induce any effect on neural differentiation

By day 50 of brain organoid development, the progenitor cells have differentiated into neuron cells, which then migrate to the outer side of the brain layer [26]. Neurons then undergo maturation that can be detected with mature neuronal markers such as NEUN, which are typically observed after 19 weeks of gestation [32]. To further assess the effect of 1.7 GHz LTE RF-EMF on the formation of layers in human brain organoids, we confirmed neuronal differentiation with the progenitor cell marker PAX6 and the neuronal marker NEUN using immunofluorescence microscopy and qPCR. In both the RF-EMF-exposed and the sham organoids, the expression of PAX6 and NEUN became clearly detectable in distinct layers as development proceeded over time (Figure 3A). These observations indicate that the progenitor cells in organoids of both groups differentiated into neurons, and the VZ and SVZ were well-segregated.

**Figure 3.**
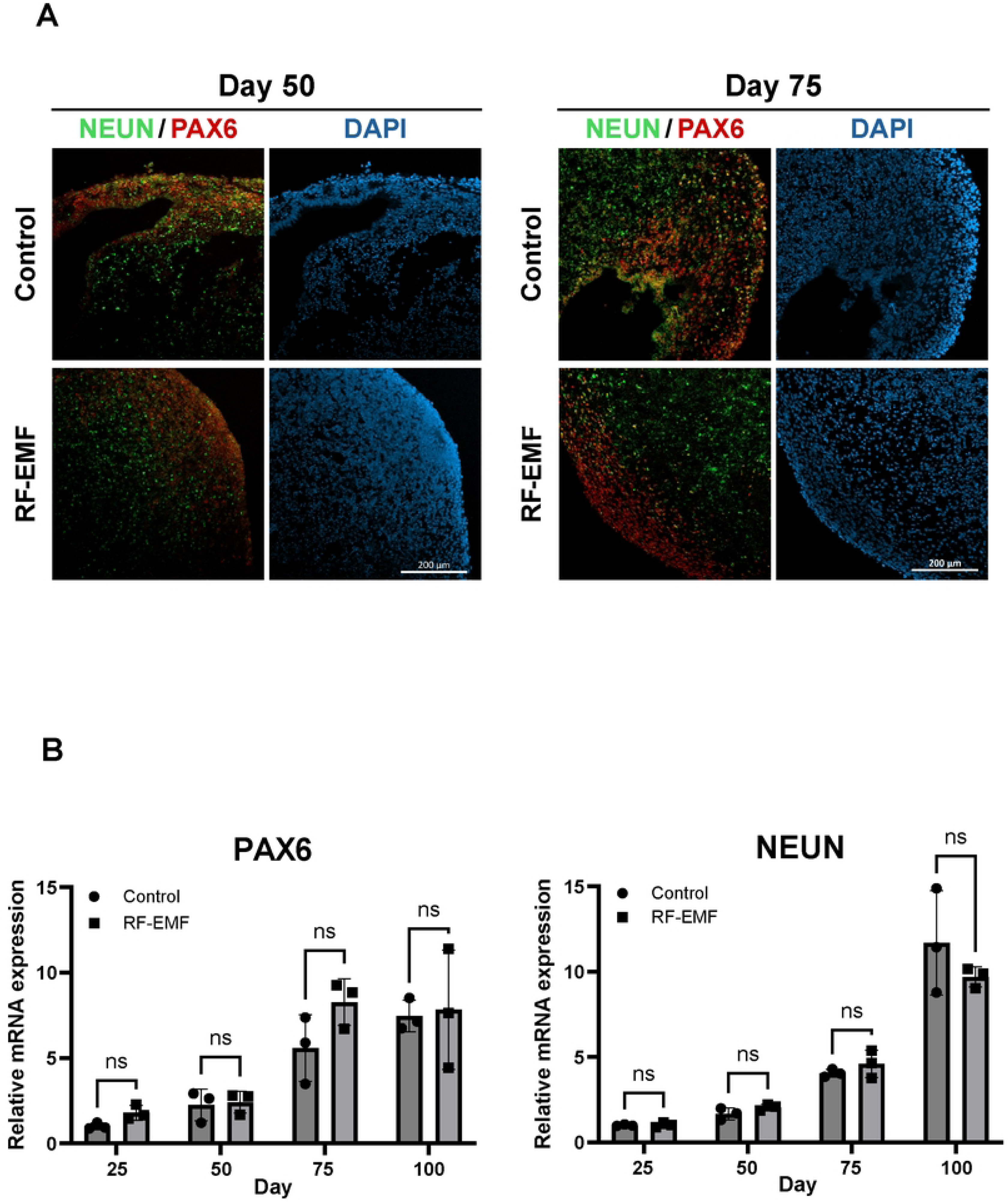
The neuronal differentiation of human cortical brain organoids exposed to 1.7 GHz LTE RF-EMF and the sham control. The developing organoids were exposed to 1.7 GHz LTE RF-EMF at a SAR of 8 W/kg for 5 h every day with the sham controls. (A) Organoids at days 50 and 75 were immune-stained with the progenitor marker PAX6 and the neuronal marker NEUN. The same confocal microscopy (LSM900, Carl Zeiss) used in the preceding figures, with 10X objectives, was employed to acquire images. Scale bar, 200 μm. (B) The expression levels of PAX6 and NEUN in developing organoids were analyzed using qRT-PCR. The expression levels of PAX6 and NEUN were all normalized using the expression of actin as a reference gene. The relative mRNA expressions of normalized target genes were plotted over the average expression of the day 25 sham control organoids. The experiments were independently replicated three times, each with 3 organoids. Multiple paired-t-tests were used to compare the expression levels of the exposed and control groups. P > 0.05 was considered statistically not significant, ns.

Moreover, when we examined the relative mRNA expression of PAX6 and NEUN during the development of brain organoids, their expressions were gradually increased over time both in the RF-EMF-exposed and the sham control organoids (Figure 3B, a slight but statistically not significant difference between the two groups). Therefore, these data suggest the possibility that exposure to 1.7 GHz LTE RF-EMF may not have an obvious effect on neuronal differentiation in human brain organoids.

### 2.5 1.7 GHz LTE RF-EMF did not indue obvious difference on synaptogenesis compared with the sham

From 17 to 22 gestational weeks, inhibitory neurons begin to connect to excitatory neurons to form neural circuits in the brain [33]. This process, called synaptogenesis, is an essential part of human brain development. If there are deficits in synaptic function, signal transmission through the neural circuit would be impaired, potentially leading to neurodegenerative diseases [34]. Thus, we investigated the risk of synapse formation in the human brain cortical organoids exposed to 1.7 GHz LTE RF-EMF at SAR for 8 W/kg daily.

We confirmed the synapse formation in the RF-EMF exposed and sham brain organoids at day 50 and 75 using the neuronal dendrite marker MAP2 and the synapse marker SYN1. Compared with the brain organoids at day 50, the day 75 organoids displayed pronounced dendrites with extended axons in both the RF-EMF-exposed and the sham organoids, when shown by immunostaining with MAP2 (Figure 4A, arrows). The number of synapses stained with SYN1 was also increased in the day 75 organoids in the two groups (Figure 4A). Consistently, the relative mRNA expression of MAP2 and SYN1 in qPCR was significantly increased between the day 50 and the day 75 organoids in both the RF-EMF-treated and the sham control, demonstrating the enhanced synapse formation in this stage of organoid development (Figure 4B). We could not detect any significant difference in synaptogenesis and its marker expression between the RF-EMF-exposed and the sham control organoids (Figure 4). When we calculated the relative increase of MAP2 and SYN1 expression in the RF-EMF-exposed and the sham between day 50 and day 75, the MAP2 expression of day 75 in the RF-EMF-exposed increased 168% of the day 50 while the sham 220%, and the SYN1 expression of day 75 in the RF-EMF-exposed increased 213% of the day 50 while the sham 288% (Figure 4B). The increased MAP2 and SYN1 expressions were a little higher in the sham compared with the RF-EMF-exposed, but their differences were statistically insignificant (Figure 4B). Altogether, these observations suggest that exposure to 1.7 GHz LTE RF-EMF at a SAR of 8 W/kg does not induce obvious detectible difference on neuronal maturation and synapse formation in the brain organoids.

**Figure 4.**
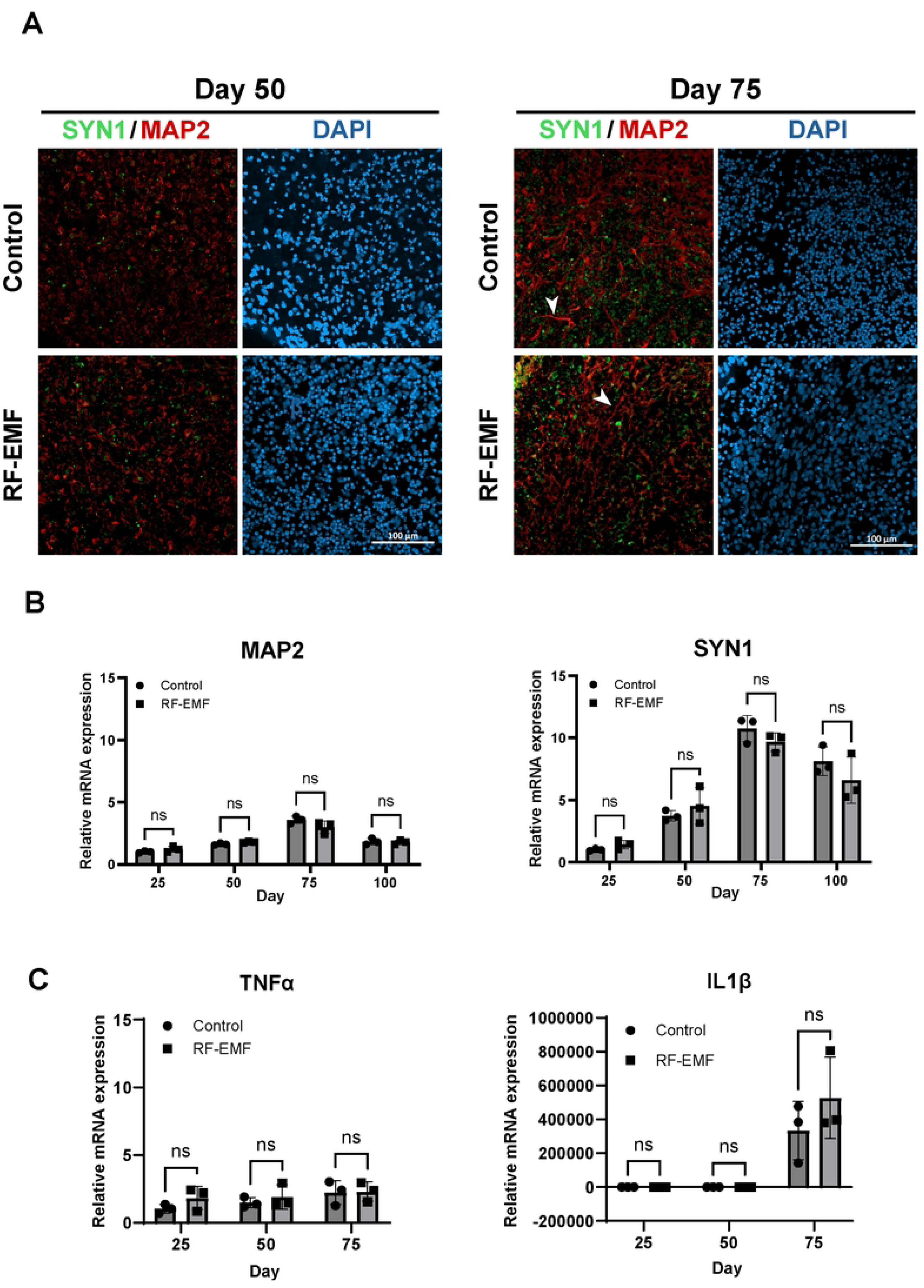
The synaptogenesis of 1.7 GHz LTE RF-EMF-exposed and the sham control organoids. The developing organoids were exposed to 1.7 GHz LTE RF-EMF at a SAR of 8 W/kg for 5 h every day with the sham controls. (A) Synaptogenesis was confirmed by staining for the synapse marker SYN1 and dendrite marker MAP2 (white arrow). The same confocal microscopy utilizing 20 X objectives was used to capture the images. Scale bar,100 μm. (B) The expression levels of (B) MAP2 and SYN1, (C) TNFα and IL-1β, in developing organoids were analyzed using qRT-PCR. The expression levels of target genes were all normalized using actin expression as reference genes. The relative mRNA expressions of normalized target genes were plotted over the average expression of the day 25 sham control organoids. The experiments were independently replicated three times, each with more than 3 organoids. Multiple paired-t-tests were used to compare the expression levels of the exposed and the control groups. P > 0.05 was considered statistically not significant, ns.

We also assessed the effect of 1.7 GHz LTE RF-EMF on the astrocyte differentiation in brain organoids by analyzing the expression of IL-1β and TNFα. According to the reported study, astrocytes begin to mature between day 50 and 100 in human cortical brain organoids [23]. Mature astrocytes secrete IL-1β and TNFα, which are essential for fetal brain development and synaptic formation. The expression of IL-1β and TNFα was measured by qPCR in both the RF-EMF-exposed and sham control human brain organoids. As reported, IL-1β expression was almost undetectable in the day 25 and 50 organoids and started to be detected in the day 75 organoids both in the RF-EMF-exposed and the sham (Figure 4C). In the day 75 organoids, we observed the difference in the IL-1β expression between the RF-EMF-treated and sham control groups, but it was statistically insignificant. Meanwhile, a low-level of TNFα was consistently detected in both the RF-exposed and sham control organoids from day 25 to day 75 (Figure 4C). The low but constant expression of TNFα in organoids at the days 25, 50, and 75 indicated that it was likely a basal-level expression and not yet induced by astrocytes.

We could not detect a statistically meaningful difference in TNFα expression between the RF-exposed and sham control (Figure 4C). These observations suggest that exposure to 1.7 GHz RF-EMF at a SAR of 8 W/kg does not significantly affect the intrinsic cytokine expression and astrocyte differentiation in human brain organoids.

## 3. Discussion

With the increasing use of wireless communication technologies, frequent exposure to RF-EMF from LTE has raised concerns regarding its potential effects on fetal development. However, there are technical challenges in designing experiments involving fetuses due to safety and ethical concerns [35, 36]. In this study, we aimed to investigate the potential risk associated with RF-EMF exposure on fetal brain development using a human brain organoid model that recapitulates the developmental processes of fetal brain.

In the previous studies, we reported that continuous exposure of various human cells to 1.7 GHz LTE RF-EMF induced decreased cell proliferation, but SAR of 0, 0.4, 1, 4, and 8 W/kg did not induce any changes in proliferation with tight thermal control [7–9]. In this study, we exposed developing human brain organoids daily for 100 days to 1.7 GHz LTE RF-EMF at SAR of 8 W/kg generated from the same device used in the previous studies. As shown in the Result (Figures 1-4), we could not observe meaningful differences in the size, neural differentiation, and the expression of markers for each developmental stage between the RF-EMF-exposed organoids and the sham controls. These observations suggested the possibility that 1.7 GHz LTE RF-EMF at a SAR of 8 W/kg may not seriously affect the progression of brain cortical organoid development. Considering that the International Commission on Non-Ionizing Radiation Protection (ICNIRP) recommended the SAR values of 4 W/kg for the limbs, 2 W/kg for the local head/torso, and 0.08 W/kg for the entire body as the safety limits for electromagnetic radiation from mobile devices for the general public [3], the strength of RF-EMF generated by mobile phones used daily would be far lower than SAR of 8 W/kg, the strength of RF-EMF we applied directly to the organoids. Additionally, a fetus is highly protected inside pregnant individuals, and its exposure would be indirect. Taken together, we expect that the physiological influence of 1.7 GHz LTE RF-EMF on early fetal brain development would be minimal. Nonetheless, our results should be confined to 1.7 GHz LTE RF-EMF, and RF-EMF with different frequencies might have different physiological outcomes on brain organoid development. To assess the precise effect of RF-EMF on human fetal brain development, further exposure studies of human cortical organoids to RF-EMF of different frequencies and a calculation of the amount of RF-EMF penetrating the body of pregnant individuals would be needed.

In this study, we could not detect meaningful difference in the images and the expressions of SOX2, PAX6, NEUN, MAP2, and SYN1 during the development of cortical brain organoids between the 1.7 GHz LTE RF-EMF-exposed groups and the sham controls (Figures 2-4 A). However, the qPCR analyses showed that the pluripotency marker PAX6 was slightly more expressed and the neuronal markers MAP2 and SYN1 were slightly less expressed in day 75 organoids of the RF-EMF-exposed group compared to the sham controls (Figures 3B, 4B). Additionally, the RF-EMF-exposed organoids at day 100 showed slightly higher expression of SOX2 and lower expression of NEUN than those of the sham control groups (Figures 2B, 3B). Thus, we cannot exclude the possibility that exposure of developing brain organoids to 1.7 GHz LTE RF-EMF may slightly delay neuronal differentiation, although all these differences were not significant in our statistical analyses. These minor differences in the timely expression of synaptogenesis markers between the RF-EMF-exposed and the sham control organoids should be further validated by other research groups with additional studies on the effect of RF-EMF on human brain organoid development. In addition, further studies on the effects of 1.7 GHz RF-EMF on various brain organoids generated with different iPSC lines should be needed to resolve the meaningful physiological outcomes of RF-EMF exposure on early brain organoid development. The main contribution of this study would be the initial application of human brain organoids generated with hiPSC as a model system for monitoring the physiological effects of RF-EMF on early fetal development including brain.

The ICNIRP safety limits for electromagnetic radiation from mobile devices for the general public are 4 W/kg for the limbs, 2 W/kg for the local head/torso, and 0.08 W/kg for the entire body. We used 8 W/kg as the SAR value of exposure in this study, which is higher than the ICNIRP limits. The reason we chose 8 W/kg as the SAR was because we observed that 1.7 GHz LTE RF-EMF at SAR of 0, 0.4, 1, 4, and 8 W/kg induced no changes in the proliferation of various human cells with the same device used in this study, and 8 W/kg is the highest SAR value without generating heat [9]. We wanted to try the strongest SAR value first because a SAR of 8W/kg has a higher chance of inducing physiological effects than other lower SAR values, if there are any effects. In this study, we showed that 1.7 GHz LTE RF-EMF at a SAR of 8 W/kg for 5 h over 100 days had no significant effect on human brain organoid development. However, to define the physiological effect of 1.7 GHz LTE RF-EMF on human organoid development more accurately, we need to examine different intensities ranging from 0.2 W/Kg to 4 W/kg in future studies.

During this study, we observed a noticeable decline in the quality of brain organoids at day 100. Both the RF-EMF-treated and sham control organoids showed reduced size and decreased expression of neuronal markers after day 75 (Figure 1-4, Figure S4). Consequently, we presented the immunofluorescence data of day 100 as supplementary in Figure S4, since the quality of the organoids was not sufficient to draw conclusions about the brain developmental processes. We speculate that as proliferation and differentiation of the neuronal cells in organoids proceed beyond a certain stage, apoptosis within the organoids may be induced due to hypoxia and nutrient deficiency [37]. Therefore, although the brain organoid serves as an excellent model to study the process and effects of environmental stresses on early human brain development, the information obtained from cortical brain organoids on later stages of development after synaptogenesis would be limited.

In conclusion, we observed that daily exposure to 1.7 GHz LTE RF-EMF at a SAR of 8 W/kg for 5 h over 100 days did not induce significant effect on human brain organoid development, suggesting a possibility that 1.7 GHz LTE RF-EMF may not cause serious damage on fetal brain development. Further studies using various human organoids from hiPSC would contribute to improving comprehension of the potential risks associated with 1.7 GHz LTE RF-EMF exposure during pregnancy.

## 4. Materials and Methods

### 4.1 IRB Permission and Maintenance of hiPSC

“Study on the effect of the daily exposure of 1.7GHz LTE electromagnetic field in fetal brain development using a human brain organoid model of the cortical cortex” obtained the permission and approval from Yonsei University Institutional Review Board (https://irb.yonsei.ac.kr). IRB Approval Number: 7001988-202404-BR-2237-01E

The hiPSC line (ASE9209) used in this study was purchased from Applied Stem Cell. ASE9209 cells of passage number 35 were cultured on Matrigel-coated 6 well cell culture plates (Corning, USA #354230), using mTeSR1 medium (STEMCELL Technology, Canada #ST85850) at 37℃ in 5% CO_2_. For passaging, hiPSCs at 70-80% confluency on the Matrigel-coated plate were washed with 1 ml of PBS (Welgene, Korea # LB 001-02) and incubated with 1 ml of gentle cell dissociation reagent (STEMCELL Technology, Canada #ST07174) for 7 min at room temperature. After removing the gentle cell dissociation reagent, 1 ml of mTeSR media was dispensed onto the cells to dissociate them into small colonies. The small colonies were subsequently distributed on new Matrigel-coated wells. ASE9209 cells in culture were monitored daily, and their stacked or differentiated colonies were removed as necessary to maintain their pluripotency. The mycoplasma infection was also monitored in ASE9209 culture by using e-Myco Valid Mycoplasma PCR detection kit (LiliF Diagnostics #25239).

### 4.2 Generation of human cortical brain organoid

To generate human cortical brain organoids, we modified the method previously published by Pasca *et al.* and Yoon *et al.* [2, 26]. hiPSCs were incubated with gentle cell dissociation reagent for 8 min at room temperature and dissociated into single cells. As an option, hiPSCs can be exposed to 1% dimethyl sulfoxide (DMSO) in mTeSR medium one day before the organoid formation to facilitate dissociation. To ensure the consistent organoid size, Aggrewell^TM^800 (STEMCELL Technology, Canada #34811) with 300 microwells was used for the formation of uniformly sized hiPSC spheroids. Between 1.5 × 10^6^ and 3.0 × 10^6^ cells were added to each well in 2 ml mTeSR medium with 10 μM Rho-kinase inhibitor Y-27632 dihydrochloride (Tocris, UK # 1254). To aggregate the cells into the microwells, the cells were centrifuged at 100 x g for 3 min and incubated in a 5% CO_2_ atmosphere at 37℃ for 24 h. The spheroids were collected by gentle pipetting up and down and placed on a 40 μm cell strainer on top of the 50 ml tube. After washing several times with PBS, the strainer was inverted on a new 50 ml tube, and hiPSC spheroids were collected by washing with 5 ml of complete medium. Spheroids were placed in the low attachment dishes (Corning, USA #4615) with Neurulation media containing Dulbecco’s modified eagle medium/ Nutrient mixture F-12 (DMEM/F12; Cytiva, USA #SH30023.01), knock-out serum replacement (Gibco, USA #10828028), 100X MEM non-essential amino acids (NEAA; Gibco, USA #11140050), Antibiotic-Antimycotic (Anti-Anti; Gibco, USA #15240-062), β-mercaptoethanol (Sigma, USA #M7522-25ml) and two SMAD pathway inhibitors, 5 μM Dorsomorphin (Sigma, USA #P5499) and 10 μM SB-431542(Tocris, UK #1614). Neurulation media supplemented with Dorsomorphin and SB-431542 was changed daily from day 1 to day 5. On the 6th day of suspension, human brain spheroids were transferred to Neural media composed of Neurobasal A medium (Gibco, USA # 271567), B27 supplement minus vitamin A (Gibco, USA # 12587010), Glutamax (Gibco, USA # 10828028), and Anti-Anti, with basic fibroblast growth factor (bFGF; R&D Systems, USA #233-FB-025) and epidermal growth factor (EGF; R&D Systems, USA # 236-EG-200). This medium was changed daily for the first 10 days and every other day until day 24. After corticogenesis, organoids were incubated in the Neural medium, where bFGF and EGF were replaced with brain-derived neurotrophic factor (BDNF; Peprotech, USA #450-02) and recombinant human neurotrophin-3 (NT-3; Peprotech, USA #450-03). The organoids were cultured in fresh medium every other day until day 42 to induce differentiation of neural progenitor cells. From day 43 onwards, they are maintained in the Neural medium without supplements, with fresh media replacement every 4 days.

### 4.3 RF-EMF device and exposure

The same 1.7 GHz LTE RF-EMF radiation system reported in the previous studies was applied for the exposure of brain organoids [7–9]. This device is a radial transmission line (RTL) exposure system, and the details of its design were described in Choi *et al.* [7]. As shown in Choi *et al.*, the SAR value was originally calculated based on temperature measurement. A dosimetric simulation of the SAR distribution on the culture dish at the exposure condition used in this study (8W/kg, 23.52 W setting) was performed by Sim4Life Tool (Zurich MedTech AG).

The RF-EMF generating device was preheated for approximately 30 min before initiating RF-EMF exposure. For RF-EMF exposure, 90 mm low-attachment dishes containing organoids were placed 13.6 cm away from the conical antenna at the core of the exposure chamber. The human brain organoids in the dishes were exposed to RF-EMF radiation from a single LTE signal operating on the Wideband Code Division Multiple Access (WCDMA) standard at 1700 MHz, with a constant SAR value of 8 W/kg for 5 h every day (Table 1). To maintain the temperature of the exposure chamber at 37 ± 0.5℃, the water circulator was consistently operated during the exposure time. The sham control group of RF-EMF-unexposed organoids was incubated under the same incubation condition as the RF-EMF chamber.

### 4.4 Cryosection

At least 3 human brain organoids grown for designated periods were fixed in 1 ml of 4% paraformaldehyde (PFA; Tech & Innovation, USA #BPP-9004) overnight at 4℃. After aspirating the 4% PFA, organoids were washed in PBS and incubated with 30% sucrose solution for 24-36 h at 4℃. The organoids were then placed in a cryo-mold with OCT compound (Leica Biosystems, Germany #3801480), frozen in liquid nitrogen, and stored at - 80℃. The cryo-block was sectioned to a thickness of 10 μm using a cryostat (Leica Biosystems, CM 1950) and attached to adhesion microscope slides (Epredia, USA #J1800AMNZ).

### 4.5 Immunofluorescence Microscopy

For immunofluorescence microscopy, cryosections were blocked in 2% goat serum (Jackson ImmunoResearch, UK #050-000-121) in 0.3% Triton X-100 for 1 h. The sections were then incubated with primary antibodies in 0.1% Triton X-100 at 4℃ overnight. After washing with PBS, they were incubated with secondary antibodies for 1 h at room temperature in dark conditions. After washing with PBS three times, the organoid sections were mounted with a mounting solution (Vector Laboratories, USA #CA94010) containing DAPI. Images of sections were captured by confocal microscopy (Carl Zeiss, LSM900) with detectors (GaAsP-PMT 2ea, additional GaAsP-PMT 1ea) and processed with ZEISS Zen version 3.9. The following primary and secondary antibodies were used with dilutions: SOX2 (1:400; Cell Signaling, USA, #3579S), PAX6 (1:100; Invitrogen, USA #42:6600), NEUN (1:500; Millipore, USA #MAB377), MAP2 (1:500; Synaptic Systems, Germany #106002), SYN1 (1:500; Synaptic Systems, Germany #188004), Alexa Fluor Rabbit 488 (Invitrogen, USA #A11008), Alexa Fluor Rabbit 594 (Invitrogen, USA #A11012), Alexa Fluor Mouse 488 (Life Technologies, USA # A11001), Alexa Fluor Guineapig 594(Invitrogen, USA #A11076).

### 4.6 Quantitative Real-time PCR (qRT-PCR)

At least 3 organoids were used for RNA extraction. RNA was extracted using an RNase-Free DNase set (Qiagen, Germany #175041095) and cDNA was synthesized by the PrimeScript^TM^RT Reagent Kit (Takara, Japan #AMG1372A). qRT-PCR was performed using a TB Green^TM^ Premix Ex Taq^TM^ (Takara, Japan #AMG1551A). The primers used are listed in Table 2.

**Table 2.**
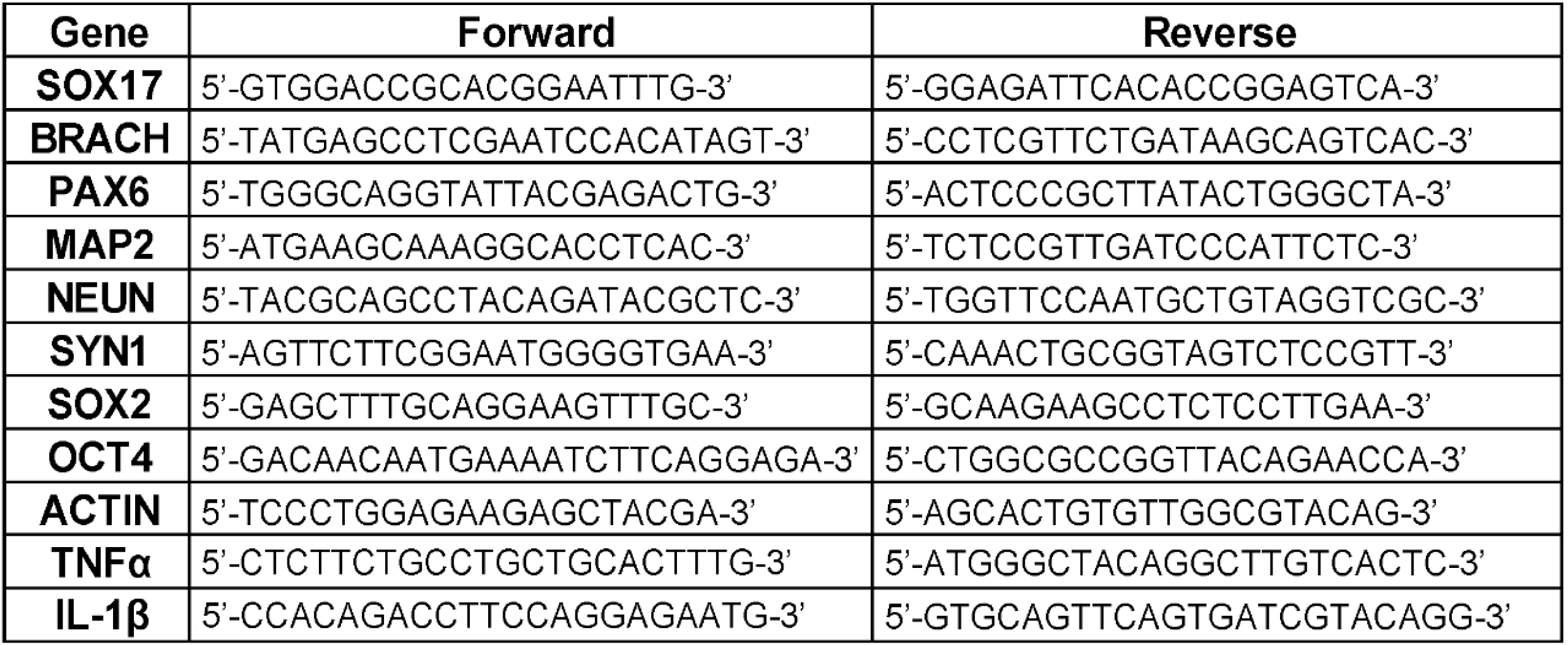
Primers used in qPCR analysis.

### 4.7 Size measurement of organoids

The size of organoids in this study was measured using the ImageJ program under identical scale settings; 0.254 distance in pixels was set as 1 micrometer in all images. Each organoid was selected using ‘Analyze particles’ or ‘Tracing tools’, and its size was measured by the ROI manager. On each designated day, the obtained average size of 10-60 organoids was normalized to the average size of day 1 organoids. Three independent experiments were performed each day, and their average was plotted.

### 4.8 Statistical analysis

All statistical analyses were conducted using GraphPad Prism version 10. Data were presented as mean ± standard deviations from three independent experiments. P-values < 0.05 (*), P < 0.01 (**), and P < 0.001 (***) were considered statistically significant, and P > 0.05 was considered statistically not significant (ns).

## 5. Acknowledgments

The authors appreciate Prof. Chul Hoon Kim (Yonsei University College of Medicine) and Dr. Baek-Soo Han (Korean Research Institute of Bioscience & Biotechnology) for technical support for the generation of human brain organoids.

## 6. Author Consent and Declaration

All authors reviewed the manuscript and consented on publication. All authors declared no conflict of interest.

## Abbreviations used

RF-EMF: radio frequency electromagnetic field
LTE: long term evolution
SAR: specific absorption rate
hiPSC: human induced pluripotent stem cell
NE: neuroepithelial cell
VZ: ventricular zone
SVZ: subventricular zone
aRGs: apical radial glial cells
RGs: radial glial cells
IPC: intermediate progenitor cell
PCW: post-conception weeks
qPCR: quantitative polymerase chain reaction

---

**Figure S1. Assessment of the quality of hiPSCs used for human cortical organoids.**

(A, B) To verify the pluripotency of hiPSCs (ASE9209), the expression of (A) OCT4 and SOX2 and (B) SOX17 (endoderm marker) and BRACH (mesoderm marker) was measured with qPCR. HeLa cells were used for expression comparison control. The expression levels of target genes in ASE9209 cells were analyzed using qPCR and normalized with actin as a reference gene. The relative mRNA expressions of normalized target genes were plotted over their expression in HeLa cells. (C) The quality of ASE9209 used in this study was verified by checking the mycoplasma infection using a PCR kit (e-Myco Valid Mycoplasma PCR detection kit, LiliF Diagnostics, #25239).

**Figure S2. The developmental progress of human cortical brain organoids at different time points.**

(A) Images of hiPSC spheroids and organoids were captured by a Nikon microscope (ECLIPSE Ts2) with a 4 X and 10 X objective. Scale bar, 100 μm (hiPSC spheroids), and 500 μm (organoids). (B) The graph represents the size of human brain organoids measured on each designated day using the ROI manager in Image J software.

**Figure S3. Proper development of internal layers including VZ and SVZ within the human cortical brain organoids.**

The expressions of (A) SOX2 (pluripotency marker), (B) PAX6 and NEUN were detected by immunofluorescence microscopy at days 25 and 50. The images were obtained using the same confocal microscopy used in Figures 2-4 with a 10X objective lens. Scale bar, 200 μm

**Figure S4. Analysis of day 100 human brain cortical organoids exposed to 1.7 GHz LTE RF-EMF daily.**

The developing organoids were exposed to 1.7 GHz LTE RF-EMF at a SAR of 8 W/kg for 5 h every day until day 100 with the sham controls. The day 100 organoids were immune-stained with (A) SOX2 (B) NEUN/PAX6, and (C) SYN1/MAP2 and detected using the confocal microscope equipped with 10 X and 20 X lenses. The white arrow illustrates the morphology of progenitor cells, and the red arrow indicates differentiation cells. Scale bars, (A) 50 μm and 200 μm, (B) 200 μm, (C) 100 μm

**Figure S5. Design of the 1.7 GHz LTE RF-EMF organoid exposure system used in this study and the evaluation of SAR values in this system.**

Details of the design of this system (a radial transmission line exposure system) were described in Choi *et al.* [7]. (A) An image of the 1.7 GHz RF-EMF exposure device. (B) The block diagram of the 1.7 GHz RF-EMF exposure system. (C) Cross-sectional view of the RTL exposure chamber. (D) Temperature measurement and linear fitting for the center point at the LTE 1.7 GHz frequency. Temperature for calculating the SAR value was measured without circulating water system to cool down during RF exposure. (E) A dosimetric simulation of the SAR distribution at 8W/kg (23.52 W setting) on the culture dish (90 mm radius with 60 ml culture medium) was performed by Sim4Life Tool (Zurich MedTech AG).

